# Concordance and dissonance: A genome-wide analysis of self-declared versus inferred ancestry in 10,250 participants from the HostSeq cohort

**DOI:** 10.1101/2025.06.10.658783

**Authors:** René L Warren, Inanc Birol, CGEn HostSeq Initiative

## Abstract

Accurate characterization of human diversity is foundational to equitable genomics. In this study, we analyzed self-declared and genome-derived ancestry in 10,250 participants from the pan-Canadian HostSeq cohort. Using the alignment-free ntRoot algorithm on whole genome sequencing data, we inferred global and local ancestry at the continental super-population level and compared these with self-reported sociocultural identity categories. We observed high concordance among individuals self-identifying as White (98.8%), Black (97.2%), East Asian (96.1%), and South Asian (89.9%). Concordance was lower among those self-identifying as Hispanic (74.6%), Middle Eastern / Central Asian (67.9%), or Indigenous (40.7%), reflecting greater admixture complexity. Agreement between expected and inferred ancestry labels was modest (Cohen’s kappa κ = −0.01 unweighted; 0.35 weighted), and ancestry discordance was strongly associated with higher Shannon entropy of ancestry fractions. Principal component analysis of ntRoot-derived ancestry composition revealed tightly clustered profiles in some groups and broader, overlapping distributions in others, illustrating how sociocultural identities and genomic data capture distinct but intersecting dimensions of human diversity. These findings support the complementary use of genome-derived continental ancestry fractions alongside self-identification, particularly in settings where sociocultural labels may be incomplete, heterogenous, or poorly aligned with genetic background. This approach can improve scientific rigor and enhance inclusion in population-scale genomics while respecting the social meaning of identity. We emphasize that genetic ancestry estimates are not proxies for race, which is a social construct with no biological basis.

## Introduction

Accurately characterizing human diversity is critical for advancing equity in biomedical research and healthcare (1). Self-declared ancestry reflects complex sociocultural, historical, and geopolitical realities, while genome-derived ancestry summarizes patterns of biological inheritance shaped by demographic history. Although both frameworks aim to describe human diversity, they often diverge in practice with implications for clinical interpretation, genetic risk assessment, and participant trust. Previous studies have documented varying degrees of concordance between self-reported and inferred ancestry, particularly in admixed populations or those shaped by colonial histories (2,3). Such dissonance can reinforce structural inequities when race and ethnicity are used uncritically as population descriptors, rather than as context-dependent, socially constructed categories informed by lived experience. At the same time, genome-based ancestry estimation has become a standard tool in population genomics, where it is often treated as a technical covariate rather than explicitly interrogated in relation to sociocultural identity (4–6).

Ancestry inference methodologies are typically classified as either global or local. Global Ancestry Inference (GAI) estimates genome-wide proportions of ancestry from broad reference groups, providing an overall summary of an individual’s genetic background. In contrast, Local Ancestry Inference (LAI) assigns ancestry to specific genomic segments by leveraging the persistence of ancestral haplotype blocks through recombination (7), thereby capturing admixture complexity at finer resolution and enabling locus-specific ancestry mapping. Recently, we introduced ntRoot, a scalable, reference-guided, alignment-free platform for both GAI and LAI (8). It leverages k-mer–based single nucleotide variant detection and integrated 1000 Genomes Project (1kGP) variant call sets to infer ancestry at the level of five continental super-population labels (AFR, AMR, EAS, EUR, SAS) (9,10).

In this study, we applied ntRoot to whole genome sequencing (WGS) data from Canada’s HostSeq initiative, a national resource comprising over 10,000 individuals affected by COVID-19 with linked clinical and self-declared ancestry data (11). Using ntRoot we inferred both global and local ancestry at the 1kGP super-population level and compared these estimates with self-declared identity categories captured through standardized questionnaires. This study represents one of the largest efforts to date to examine concordance and discordance between sociocultural identity labels and super-population-level ancestry predictions in a Canadian cohort, providing a quantitative view of where these dimensions converge and where they diverge within a multicultural context. We emphasize that the ancestry labels used by ntRoot are geographically defined super-population categories derived from reference panels and are not proxies for race, which is a social construct without biological basis. Our use of genomic ancestry is intendeds to complement, not replace self-identification in efforts to improve representational equity and scientific validity in genomics research.

## Methods

### Cohort Description

We analyzed data from 10,250 participants enrolled in the HostSeq project, representing a diverse cross-section of the Canadian population. Participants provided self-declared ancestry through structured questionnaires, selecting from categories aligned with national demographic standards. We acknowledge that pandemic-focused enrollment may have influenced participant diversity. For clarity, the total cohort comprises all 10,250 participants with available genomic data. Analyses requiring a direct comparison between self-declared identity and genome-derived global ancestry were restricted to an analyzed HostSeq cohort subset, defined as participants who provided a self-declared category that could be mapped to a continental-level expected ancestry (Table 1).

**Table 1.** Concordance rate by self-declared ancestry category. Concordance is defined as the proportion of individuals whose genome-derived global ancestry (GAI, assigned based on their largest local ancestry fraction) matches the expected ancestry for their self-declared identity category. The five continental-level population groups include AFR (African), AMR (Admixed American), EAS (East Asian), EUR (European), and SAS (South Asian). The total number of participants (n) is shown for each self-declared category; together, these counts represent the ‘total cohort’. Only individuals with an assigned GAI label represent the ‘analyzed HostSeq cohort subset’, as defined in Methods.

| Self-Declared Category | GAI | n | Concordant (n) | Concordance Rate |
| --- | --- | --- | --- | --- |
| Ashkenazi Jewish | EUR | 62 | 62 | 1.000 |
| Sephardic Jewish | EUR | 2 | 2 | 1.000 |
| White | EUR | 2,743 | 2,709 | 0.988 |
| Black | AFR | 213 | 207 | 0.972 |
| East Asian / Pacific Islander | EAS | 358 | 344 | 0.961 |
| South Asian | SAS | 358 | 322 | 0.899 |
| Hispanic | AMR | 185 | 138 | 0.746 |
| Middle Eastern or Central Asian | EUR | 162 | 110 | 0.679 |
| Indigenous (First Nations, Métis, Inuit) | AMR | 27 | 11 | 0.407 |
| Not Disclosed | NA | 4,723 | NA | NA |
| Prefer not to answer / Do not know | NA | 1,168 | NA | NA |
| More than one race | NA | 171 | NA | NA |
| Other | NA | 78 | NA | NA |
| **Total** | NA | **10,250** | NA | NA |

### Genetic Ancestry Inference

We used the ntRoot algorithm (v1.1.5) to perform ancestry prediction from WGS data, categorizing individuals into five continental-level population groups: AFR (African), AMR (Admixed American), EAS (East Asian), EUR (European), and SAS (South Asian). The ntRoot method employs a reference-guided framework optimized for WGS, targeting high-resolution classification. First, each individual read set in CRAM format was converted to fastq using the command: samtools fastq -@ 48 --reference GRCh38_full_analysis_set_plus_decoy_hla.fa −1 SR.1.fastq.gz −2 SR.2.fastq.gz sample.cram. Next, we executed ntRoot (v1.1.5, with parameters - k 55 --draft GRCh38.fa.gz --reads SR -t 48 -Y 0.55 −l 1000GP_integrated_snv_v2a_27022019.GRCh38.phased_gt1.vcf.gz --lai). A total of 10 batched jobs, each containing instructions to process 1,025 samples, ran for a total of 2 months and 3 weeks on server-class systems with Intel(R) Xeon(R) CPU E7-8867 v3 @ 2.50GHz and 2.6TB RAM.

### Concordance Rate Estimation

We defined an expected global ancestry for each self-declared category based on Canadian census categories, prior literature on population structure in these groups, and 1kGP continental label conventions (e.g., “Ashkenazi Jewish” → EUR, “East Asian / Pacific Islander” → EAS, etc.) (Table 1) (12–14). These mappings are heuristic and intended to reflect broad, historically grounded expectations rather than precise genetic truths. Concordance was defined as a match between the expected global ancestry and the ancestry inferred from ntRoot, assigned based on the largest predicted LAI fraction. All data handling and analysis were performed using R (v4.3.1) with the dplyr (v1.1.2) package.

### Cohen’s Kappa Agreement Analysis

To evaluate categorical agreement beyond chance, we used the cohen.kappa() function from the psych package (v2.3.9) on a contingency table comparing expected versus inferred ancestry labels (alpha level = 0.05). We reported both unweighted and weighted kappa scores, the latter accounting for partial similarity across related continental superpopulation groups.

### Shannon Entropy (Admixture Complexity)

To assess genomic admixture per individual, we calculated Shannon entropy from continental ancestry fractions (AFR, AMR, EAS, EUR, SAS) using the entropy package (v1.3.1). Entropy scores were then aggregated by self-declared category to identify which groups exhibit more mixed ancestry profiles. In this five-ancestry framework, entropy near 0 reflects effective ‘single-ancestry’ profiles, while entropy values above 1.5 correspond to substantial contributions from at least three continental ancestries.

### Principal Component Analysis (PCA)

We conducted PCA on the ntRoot-predicted ancestry fractions to visualize similarities and differences in genomic ancestry composition across self-declared identity groups. Principal components were plotted and colored by self-declared category to observe clustering and spread. PCA was computed using the base prcomp() function in R, and visualized using ggplot2 (v3.4.0).

### Inclusion/Exclusion in Analyses

Individuals selecting ‘Prefer not to answer / do not know’ were retained for completeness in descriptive summaries but excluded from concordance, entropy, and PCA analyses, as their self-identification could not be meaningfully compared to an expected ancestry label.

## Results

The HostSeq cohort comprises 10,250 individuals recruited across Canada through COVID-19 pandemic-related studies. While the largest proportion of participants (26.8%) self-identified as White (n = 2,743), smaller proportions identified as South Asian (n = 358; 3.5%), East Asian / Pacific Islander (n = 358; 3.5%), Black (n = 213; 2.1%), Hispanic (n = 185; 1.8%), Middle Eastern or Central Asian (n = 162; 1.6%), and Indigenous (First Nations, Métis, Inuit; n = 27; 0.3%). A notable fraction of individuals did not disclose their ancestry (n = 4,723; 46.1%), or selected “Prefer not to answer / do not know” (n = 1,168; 11.4%), “More than one race” (n = 171; 1.7%), or “Other” (n = 78; 0.8%) (Table 1). These uncategorized responses, which together represented the largest fraction (i.e., ∼60%) of individuals, mapped across diverse ancestries, suggesting complex identity constructs or reluctance to engage with constrained identity categories, highlighting the limits of categorical identity labels. This composition does not necessarily mirror the broader demographic diversity of Canada and likely reflects a combination of cohort recruitment patterns, barriers to participation among some communities, and the constrained nature of categorical identity labels used in biomedical research. Participants who selected ‘Prefer not to answer’ were retained in the dataset for completeness but excluded from identity-discordance calculations and ancestry visualizations, as described in Methods.

### Patterns of Discordance

Substantial discordance between expected and inferred continental labels was observed in certain categories, including individuals of self-declared Hispanic, Indigenous and Middle Eastern / Central Asian backgrounds. Among participants identifying as Hispanic, most were assigned global ancestry labels of AMR (74.6%), EUR (15.1%), or AFR (8.6%) (Fig. 1), and showed appreciable local ancestry contributions from these same groups (Fig. 2). Many individuals identifying as First Nations, Métis, or Inuit clustered within either the AMR (40.7%), or EUR (59.3%) GAI predictions. For Middle Eastern or Central Asian, a similar dispersion across EUR (67.9%), SAS (19.8%), and AFR (12.3%) continental labels suggests diverse ancestral backgrounds and limited resolution of these identities within current 1kGP-based reference panels (Fig. 1A and 1B).

**Fig. 1.**
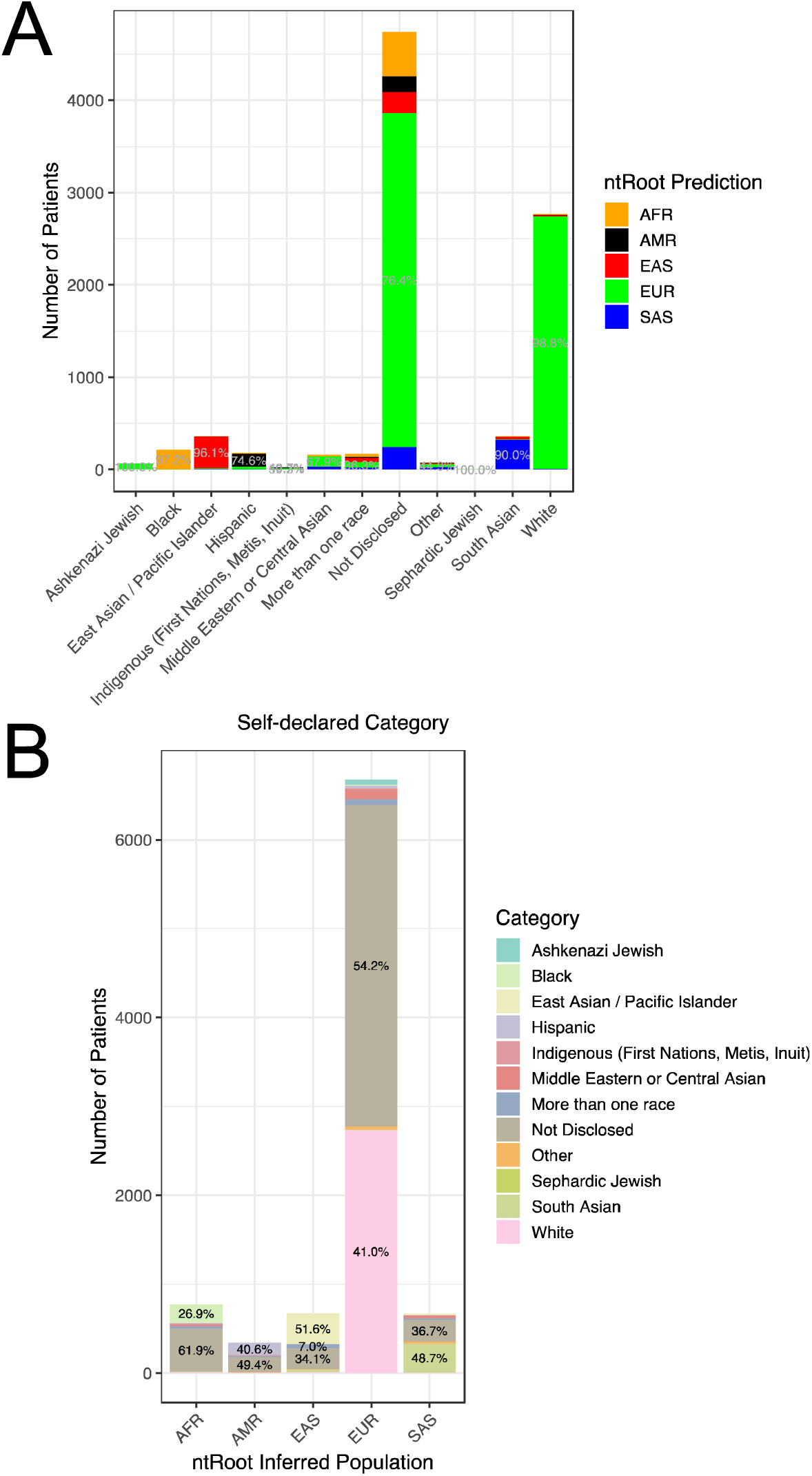
Inferred versus self-declared ancestry in the HostSeq cohort. **A)** by ntRoot predictions of the main continental ancestry (1000 Genomes Project continental ancestry labels (12) AMR:American; AFR:African; EUR:European; SAS:South Asian; EAS:East Asian. Percentages 30% or higher indicated) and **B)** by self-declared ancestry categories in the HostSeq study (percentages above 5% indicated). Participants who selected ‘Prefer not to answer’ were excluded to avoid visual stratification or interpretation of inferred ancestry in relation to an explicitly withheld identity label.

**Fig. 2.**
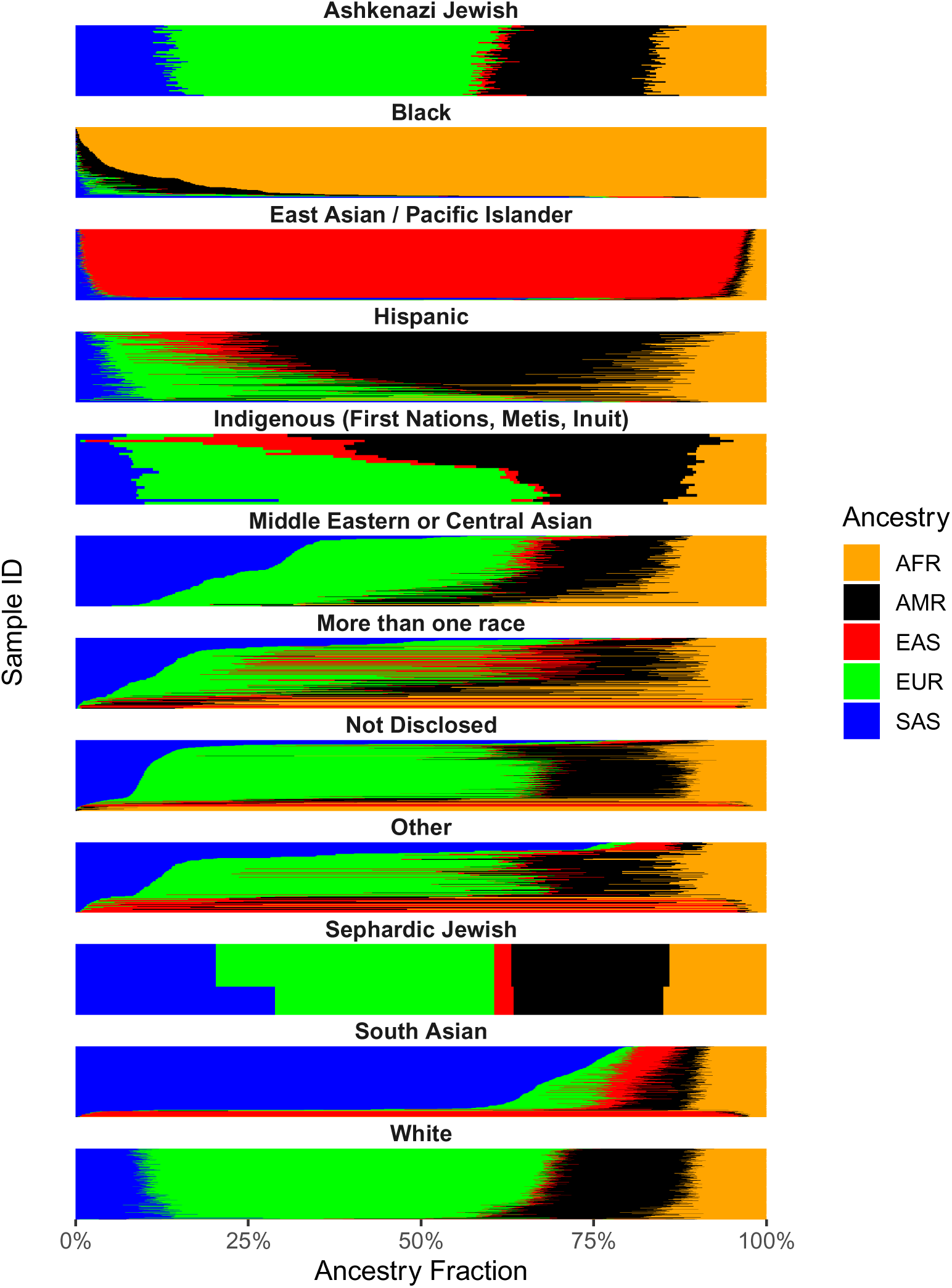
ntRoot-predicted ancestry fractions based on LAI. (x-axis, 1000 Genomes Project continental ancestry labels (12) AMR:American; AFR:African; EUR:European; SAS:South Asian; EAS:East Asian), for each HostSeq individual sample (y-axis) in the cohort, organized by self-declared identity category and normalized by sample count. Participants who selected ‘Prefer not to answer’ were excluded to avoid visual stratification or interpretation of inferred ancestry in relation to an explicitly withheld identity label.

Overall, unweighted Cohen’s kappa comparing expected and inferred continental labels was –0.01, indicating minimal agreement beyond chance across all categories (Table 2, Supplementary Fig. 1), whereas weighted kappa was 0.35, reflecting moderate agreement when accounting for partial similarity across ancestry types. Shannon entropy, a measure of admixture complexity where higher values reflect greater ancestry heterogeneity, revealed marked differences across groups. Individuals identifying as Sephardic (mean entropy [m.e.]=2.04) and Ashkenazi Jewish (m.e.=1.91), Middle Eastern or Central Asian (m.e.=2.00), Indigenous (m.e.=1.78), White (m.e.=1.72), Hispanic (m.e.=1.71), and “More than one race” (m.e.=1.72) exhibited the highest levels of admixture, consistent with ancestry contributions from multiple continental groups in the ntRoot-derived fractions, whereas South Asian (m.e.=1.36), Black (m.e.=0.474), and East Asian / Pacific Islander (m.e.=0.471) showed the lowest entropy values (Table 3, Fig. 2). Principal component analysis of ntRoot-derived ancestry fractions revealed distinct clustering among individuals self-identifying as White, East Asian, South Asian, and Black (Supplementary Fig. 2), consistent with lower admixture entropy and high concordance with inferred ancestry. In contrast, individuals identifying as Indigenous, Hispanic, or Middle Eastern / Central Asian exhibited broader, overlapping distributions in PCA space, reflecting more complex ancestry profiles and lower overall concordance between self-declared and genome-derived ancestry. These observations are consistent with the concordance table, which shows higher match rates for White (98.8%), Black (97.2%), East Asian / Pacific Islander (96.1%), and South Asian (89.9%) individuals, and lower rates for Hispanic (74.6%), Middle Eastern or Central Asian (67.9%) and Indigenous (40.7%) populations, according to their respective assigned global ancestry (Table 1).

**Table 2.** Cohen’s Kappa agreement (Interrater agreement). Analyses were performed on the analyzed HostSeq cohort subset, as defined in Methods. The five continental-level population groups include AFR (African), AMR (Admixed American), EAS (East Asian), EUR (European), and SAS (South Asian).

| **Inferred vs. Expected** | **AFR** | **AMR** | **EAS** | **EUR** | **SAS** |
| --- | --- | --- | --- | --- | --- |
| **AFR** | 1.000 | 0.500 | 0.015 | 0.400 | 0.250 |
| **AMR** | 0.000 | 1.000 | 0.175 | 0.688 | 0.100 |
| **EAS** | 0.000 | 0.000 | 1.000 | 0.312 | 0.710 |
| **EUR** | 0.000 | 0.000 | 0.000 | 1.000 | 0.390 |
| **SAS** | -0.029 | 0.000 | -0.029 | -0.059 | 1.000 |

**Table 3.**
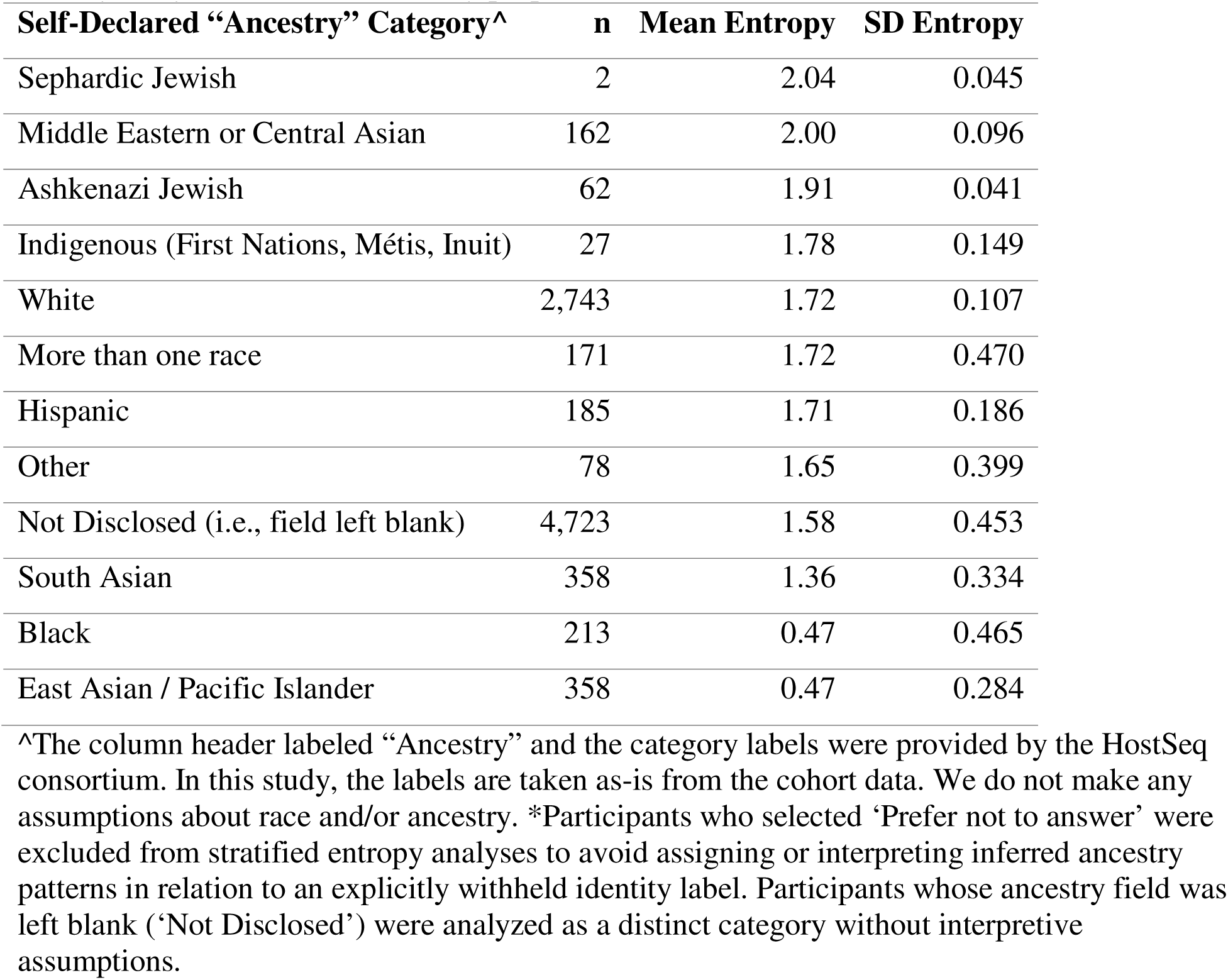
Shannon entropy and admixture by self-declared ancestry category for the HostSeq cohort, excluding participants who selected “Prefer not to answer”*. Mean entropy represents the average genomic admixture complexity for individuals within each self-declared category, based on the distribution of genome-wide ancestry fractions. Standard deviation (SD) reflects the variability of entropy within each group. Higher entropy indicates greater heterogeneity in continental ancestry proportions, and thus more admixture.

| **Self-Declared “Ancestry” Category<sup>^</sup>** | **n** | **Mean Entropy** | **SD Entropy** |
| --- | --- | --- | --- |
| Sephardic Jewish | 2 | 2.04 | 0.045 |
| Middle Eastern or Central Asian | 162 | 2.00 | 0.096 |
| Ashkenazi Jewish | 62 | 1.91 | 0.041 |
| Indigenous (First Nations, Métis, Inuit) | 27 | 1.78 | 0.149 |
| White | 2,743 | 1.72 | 0.107 |
| More than one race | 171 | 1.72 | 0.470 |
| Hispanic | 185 | 1.71 | 0.186 |
| Other | 78 | 1.65 | 0.399 |
| Not Disclosed (i.e., field left blank) | 4,723 | 1.58 | 0.453 |
| South Asian | 358 | 1.36 | 0.334 |
| Black | 213 | 0.47 | 0.465 |
| East Asian / Pacific Islander | 358 | 0.47 | 0.284 |

<sup>^</sup>The column header labeled “Ancestry” and the category labels were provided by the HostSeq consortium. In this study, the labels are taken as-is from the cohort data. We do not make any assumptions about race and/or ancestry. \*Participants who selected ‘Prefer not to answer’ were excluded from stratified entropy analyses to avoid assigning or interpreting inferred ancestry patterns in relation to an explicitly withheld identity label. Participants whose ancestry field was left blank (‘Not Disclosed’) were analyzed as a distinct category without interpretive assumptions.

### Proportional Ancestry Patterns

Figure 2 and Supplementary Fig. 3 present the ancestry composition of each self-declared group using stacked ancestry fractions. In Supplementary Figure 3, all individuals are shown in full cohort proportions whereas Fig. 2 normalizes by sample count per group to eliminate size bias. In those figures, the ancestry fractions tallied from LAI for each sample reveal that while categorical self-declared groups including Black, South Asian, and East Asian display tightly clustered and more homogenous genetic profiles, groups such as Hispanic, Indigenous, Ashkenazi and Sephardic Jewish, Middle Eastern and White contain a wider range and appreciable fractions of continental ancestries. This indicates that admixture is a common feature of these identities in this cohort, rather than an artifact of sample size or distribution.

## Discussion

Despite relatively high overall concordance between self-declared identity categories and genome-derived global ancestry predictions, entropy calculations and local ancestry fractions revealed high admixture complexity in several groups. These included individuals identifying as Ashkenazi and Sephardic Jewish, Middle Eastern or Central Asian, Indigenous, White, and Hispanic, as well as those selecting “More than one race”, “Other”, or who did not self-declare ancestry, mirroring the broad clustering patterns observed in PCA. These results reinforce the association between entropy, discordance, and generally less distinct PCA clustering in groups with more complex admixture histories.

The analysis presented here affirms the general reliability of self-declared ancestry in broad continental contexts, while also highlighting the need for more nuanced interpretation in admixed or historically complex populations. Our findings are consistent with previous work (2,15) showing that discordance between self-identified and inferred ancestry is most pronounced in groups shaped by colonial histories or recent admixture. Importantly, we emphasize that neither genome-derived ancestry nor self-identification should be treated as inherently more valid. Each reflects a different lens on identity, biological and sociocultural, and their intersection offers a richer and more complete picture of human diversity.

For genomic research, these findings reinforce the need for flexible, multidimensional approaches to participant categorization, and call for a deliberate commitment to equity-informed study design (4). The inclusion of proportional ancestry analysis underscores that identity categories such as Ashkenazi and Sephardic Jewish, Middle Eastern or Central Asian, Indigenous, White and Hispanic are not genetically homogeneous groups but socially meaningful designations encompassing diverse ancestry profiles. These insights have critical implications for how populations are stratified in biomedical research, particularly in precision medicine frameworks that rely heavily on genetic clustering and inference.

Moreover, the incorporation of local ancestry inference (LAI) using whole genome sequencing (WGS) via ntRoot provides resolution that global ancestry inference (GAI) alone cannot achieve. While GAI summarizes continental-scale proportions, LAI identifies the likely super-population origin of specific genomic segments, enabling locus-level insights into admixture and genomic architecture. This is particularly valuable in large cohort association studies, pharmacogenomics, and polygenic risk score development, especially in admixed or underrepresented populations (16–18). Fine-grained ancestry analysis at the continental level helps mitigate biases introduced by majority-European training datasets (19) and allows for the identification of ancestry-specific associations with complex traits. As such, LAI represents an essential complement to GAI in efforts to make genomic medicine more equitable and mechanistically precise.

This study also has several limitations. First, it relies on categorical mapping assumptions that assign self-declared identities to expected genomic ancestry groups, which may oversimplify complex sociocultural constructs. Second, while ntRoot offers both global and local inference, its resolution is currently limited to continental-scale super-population labels defined using 1kGP reference data and may obscure the continuous and temporally deep nature of human population structure (8,20). As others have noted, these classifications can conflate distinct concepts such as race, ethnicity, geography, and genetic ancestry (21–24), which may limit their interpretability, particularly in diverse or historically admixed populations. Additionally, the self-declared data reflect heterogeneous interpretations of ancestry, race, and ethnicity by participants. Because individuals who did not provide a specific ancestry label constituted a large fraction of the cohort, concordance and entropy estimates were necessarily restricted to the subset who did self-identify. These results should therefore be interpreted as characterizing patterns among declarative respondents rather than the entire HostSeq cohort. However, comparison of inferred global ancestry between responders and non-responders revealed statistically detectable differences driven by the large sample size, with small absolute shifts in ancestry proportions (typically <3%; data not shown). Systematic differences in identity disclosure therefore cannot be excluded. Finally, reference panels used for ancestry inference may underrepresent certain populations, particularly Middle Eastern and Indigenous groups, potentially reducing accuracy in these categories (25,26). It is important to note that the Indigenous category in this Canadian cohort may encompass a range of identities, including First Nations, Inuit, and particularly Métis individuals, whose genetic ancestry often reflects historical admixture with European populations, which may partly explain our observations. These findings, therefore, may not generalize to Indigenous populations in other national contexts, and highlight the importance of region-specific considerations, including documented colonial and post-colonial histories, when interpreting genomic diversity within socially defined groups (27–29).

## Conclusion

This work represents a large-scale effort to compare self-declared and genome-derived ancestry predictions in a nationally diverse cohort. While self-declared identity may reflect lived experience and social determinants, ntRoot provides standardized whole genome–based estimates of genomic ancestry, with advantages in scalability, reproducibility, and population-level harmonization. By combining whole genome-based ancestry inference with participant-reported identity data, we provide a picture of where genomic and sociocultural understandings of ancestry converge and where they diverge. These findings underscore the importance of integrating both biological and self-identified dimensions of identity in genomic research. Doing so will be critical for developing more inclusive, precise, and equitable research in population genomics.

## Author Contributions

Study concept: RLW, IB. Software implementation: RLW. Data analysis: RLW.

Manuscript development: RLW. Manuscript editing: RLW, IB. Funding acquisition: IB, RLW.

## Data and Code Availability

The data used in this study is available for download at CGEn (https://www.cgen.ca/hostseq-application-form). ntRoot is freely available on GitHub (https://github.com/bcgsc/ntroot).

