## Supplementary Material for "Concordance and dissonance: A genome-wide analysis of self-declared versus inferred ancestry in 10,250 participants from the HostSeq cohort"

**Supplementary Figures**

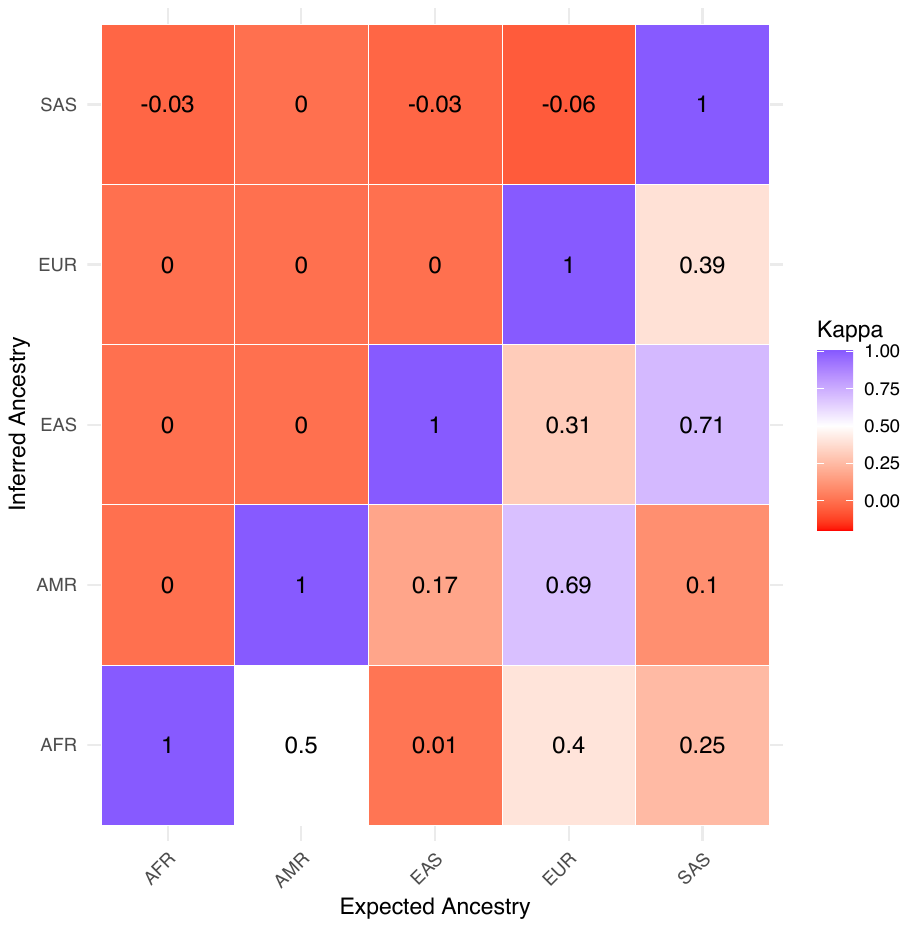

**Supplementary Fig. 1. Concordance matrix of self-declared vs. inferred ancestry.** Analyses were performed on the analyzed HostSeq cohort subset, as defined in Methods. Heatmap representing pairwise Cohen’s Kappa agreement statistics between expected and inferred continental ancestry groups. Higher agreement is shown in blue, while lower or negative agreement is shown in red. Diagonal elements represent perfect agreement, and off-diagonal values reflect partial overlap or misclassification.

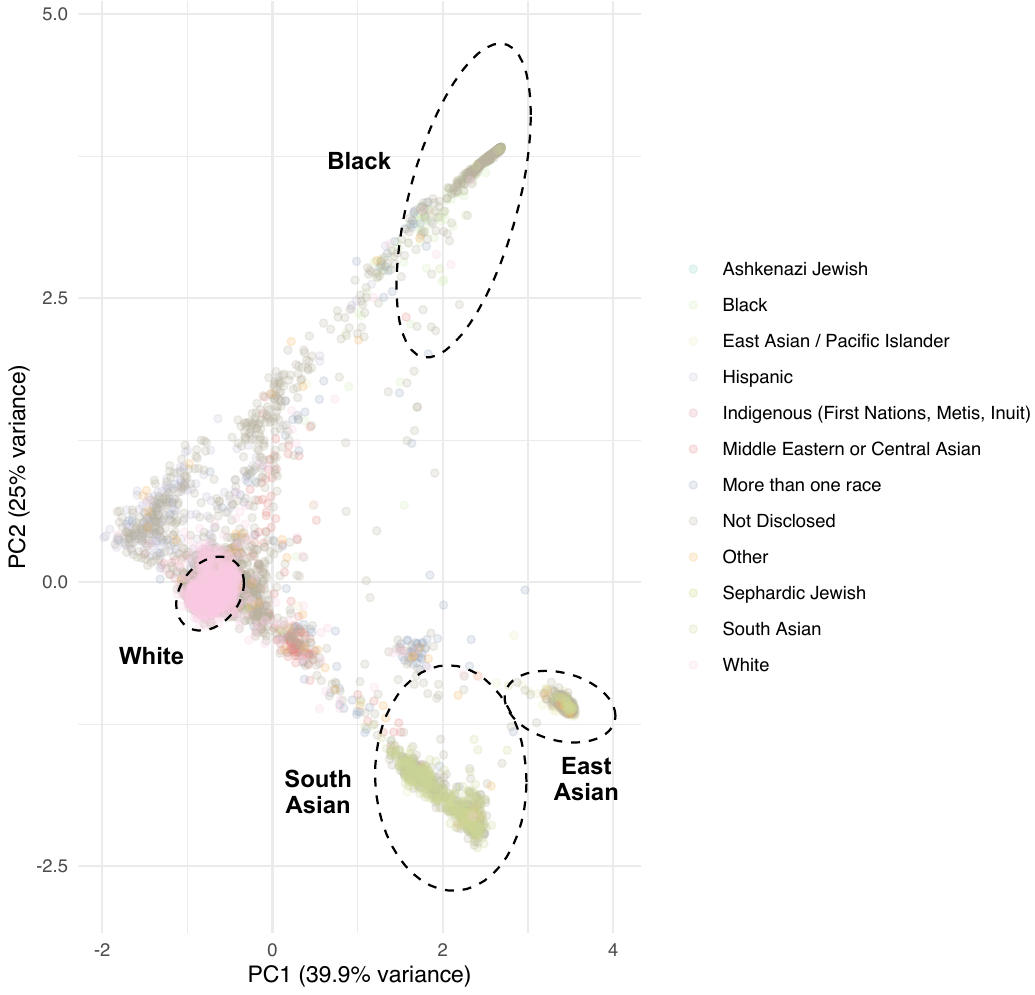

**Supplementary Fig. 2. PCA of ntRoot-predicted ancestry fractions (1000 Genomes Project continental ancestry labels AMR:American; AFR:African; EUR:European; SAS:South Asian; EAS:East Asian) across the HostSeq cohort.** Each point represents an individual, colored by their self-declared ancestry. Clustering is observed for self-identified White, Black, East Asian, and South Asian participants, while greater spread is observed among Hispanic, Indigenous, and Middle Eastern/Central Asian individuals, consistent with higher admixture complexity. Participants who selected ‘Prefer not to answer’ were excluded to avoid visual stratification or interpretation of inferred ancestry in relation to an explicitly withheld identity label.

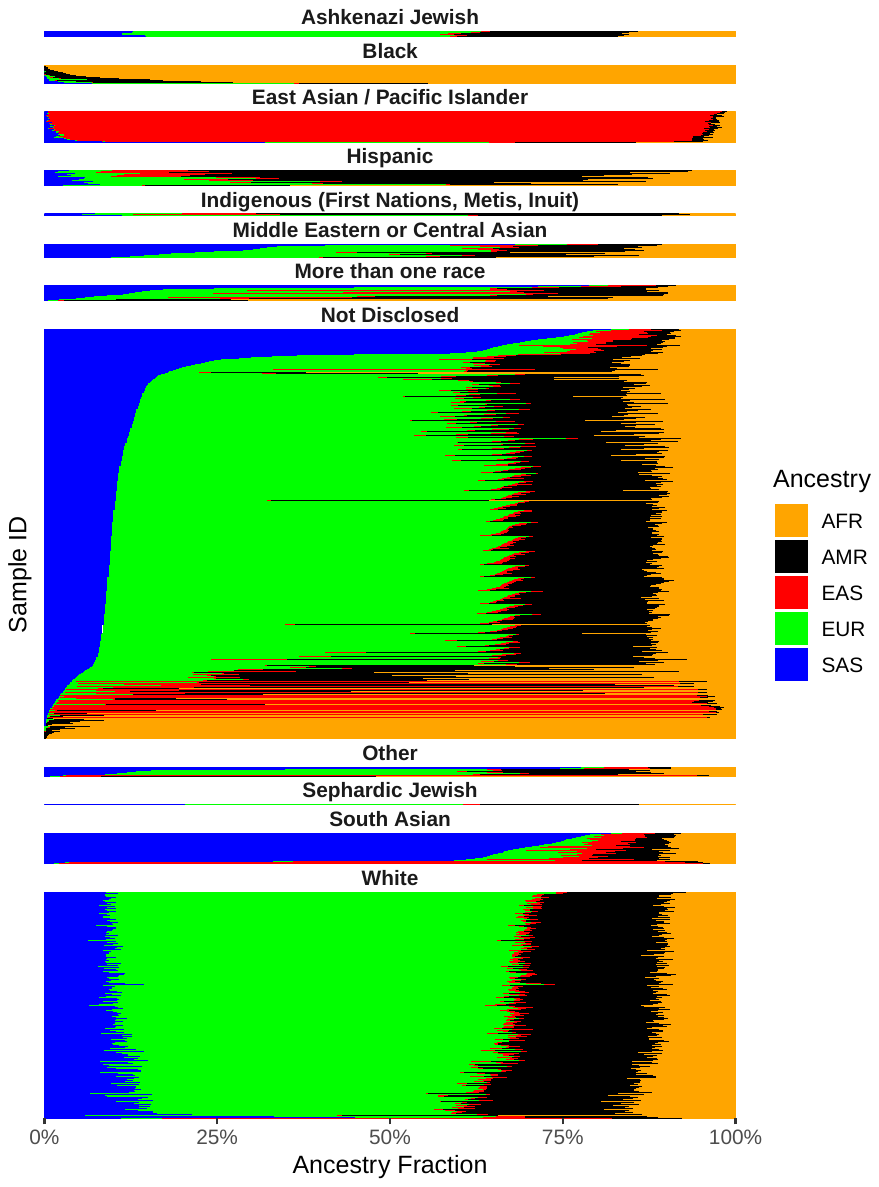

**Supplementary Fig. 3.** **ntRoot-predicted ancestry fractions based on LAI** (x-axis, 1000 Genomes Project continental ancestry labels AMR:American; AFR:African; EUR:European; SAS:South Asian; EAS:East Asian), for each HostSeq individual sample (y-axis), organized by self-declared identity category and showing full cohort proportions. Participants who selected ‘Prefer not to answer’ were excluded to avoid visual stratification or interpretation of inferred ancestry in relation to an explicitly withheld identity label.
